# Scene-selective regions encode the vertical position of navigationally relevant information in young and older adulthood

**DOI:** 10.1101/2023.10.18.562731

**Authors:** Marion Durteste, Luca R. Liebi, Emma Sapoval, Alexandre Delaux, Angelo Arleo, Stephen Ramanoël

## Abstract

Position within the environment influences the navigational relevance of objects. However, the possibility that vertical position represents a central object property has yet to be explored. Considering that the upper and lower visual fields afford distinct types of visual cues and that scene-selective regions exhibit retinotopic biases, it is of interest to elucidate whether the vertical location of visual information modulates neural activity in these high-level visual areas. The occipital place area (OPA), parahippocampal place area (PPA) and medial place area (MPA) demonstrate biases for the contralateral lower visual field, contralateral upper visual field, and contralateral hemifield, respectively. Interesting insights could also be gained from studying older adulthood as recent work points towards an age-related preference for the lower visual field. In the present study, young and older participants learned the position of a goal in a virtual environment that manipulated two variables: the vertical position of navigationally-relevant objects and the presence of non-relevant objects. Results revealed that all three scene-selective regions parsed the vertical position of useful objects independently of their subtending retinotopic biases. It therefore appears that representations in the higher-level visual system combined information about vertical position and navigational value for wayfinding purposes. This property was maintained in healthy aging emphasizing the enduring significance of visual processing along the vertical dimension for spatial navigation abilities across the lifespan.

## Introduction

Successful wayfinding in humans is dependent upon intricate processing to perceive and interpret available visual stimuli in a spatially meaningful way. Indeed, visual cues such as objects can act as anchoring points to enable the encoding of spatial relations and the formation of unified mental representations of the environment^1^. Among the multitude of properties that influence the navigational relevance of visual information, location in space has garnered much attention^2^. Behavioral and neuroimaging studies have shown that objects at positions that require a decision to be made, such as intersections, are of considerable importance for adequate navigation performance^3,4^. Proximity of objects to a boundary or to a target location also tend to facilitate the use of visual information for wayfinding purposes^5–7^. In addition, while landmarks situated in near-distance (*i.e.,* proximal space) to the observer allow for immediate visually guided navigation, those in far-distance (*i.e.,* distal space) enable map-like orientation^8–10^. Despite a wealth of research emphasizing the pertinence of object location for navigation, one critical aspect that remains uncharted is position within the vertical space.

The vertical position of information for spatial navigation is worthy of interest for three reasons. First, visual information is not processed homogeneously across the visual field. The human upper and lower visual fields exhibit striking functional disparities as the former allows for enhanced visual search and object identification^11–13^ and the latter favors acute visual acuity, contrast sensitivity and hue discrimination^14–16^. These vertical asymmetries have ramifications on cognitive abilities that are essential to spatial navigation. For example, performance on item relocation and memory tasks is improved when stimuli are presented in the lower visual field^17,18^. Second, visual cues within the upper and lower visual fields naturally carry distinct functions for wayfinding. The upper visual field is primarily associated with stable and immovable objects such as monuments or large advertising signs. The latter make up the building blocks for the creation of mental maps. The lower visual field encloses small moving objects and navigational affordance information, the processing of which is indispensable to avoid obstacles and to guide immediate movements^19–21^. Third, recent work uncovered that visual field maps are ubiquitous throughout the brain^22–24^. They were first identified in early visual areas with V1 exhibiting an inverted contralateral visual field representation and ventral and dorsal V2 and V3 areas comprising representations of the upper or lower contralateral visual field respectively^25–27^. Such visual field maps extend from the early visual cortex to key regions of the human spatial navigation network including the occipital place area (OPA), parahippocampal place area (PPA), medial place area (MPA) and hippocampus^28–30^. The OPA, PPA and MPA are ventromedial posterior regions that shape the human scene-selective network. All three regions are broadly involved in scene perception, spatial navigation and the processing of visual cues, but they each harbor specific functions. The OPA is situated on the dorsal stream, and it processes visual information that serves immediate navigational purposes such as obstacles, affordances, boundaries and egocentric distance^31–35^. Moreover, the OPA displays a contralateral lower retinotopic bias^28^. The PPA is located on the ventral stream, it is selectively engaged during scene categorization and represents the summary statistics of scene content^36–39^. In contrast to the OPA, the PPA exhibits a retinotopic bias for information in the upper visual field^28^. Finally, the MPA (or RSC)^29^ within the medial parietal cortex subserves the formation of map-based representations by encoding information such as facing direction and permanent landmark location^40–42^. Akin to the hippocampus, the MPA boasts a more general contralateral bias without any documented vertical preference^29,30^. A missing piece of the puzzle pertains to understanding whether the roles of the underlying retinotopic preferences in the OPA, PPA and MPA interact with scene processing and task demands for spatial navigation. Indeed, if retinotopic biases have direct behavioral correlates, they should not only encode task-relevant information but also task-irrelevant information. Therefore, we wondered whether spatial representations of the environment could factor in information about the vertical position of visual cues within the visual field, and whether the coding of vertical position is contingent upon navigational relevance.

The above interrogations are of particular interest in older adulthood. New results are emerging to suggest that healthy aging is associated with an increased reliance on information located in the lower visual field. Target detection, visual search and spatial memory are better performed when stimuli are presented in the lower visual field than in the upper visual field of older adults^18,43,44^. Moreover, older individuals show a systematic downward gaze bias during navigation in real and virtual spaces^45,46^. Interestingly, recent neuroimaging evidence hints at the notion that age-related differences in spatial navigation could stem partly from disparities in the perceptual computations associated with landmark processing^7,47,48^. For example, a fMRI study found that older participants recruited the OPA to a greater extent than younger participants during a landmark-based navigation task^48^. These results stress the intriguing possibility that OPA activity in aging might reflect the age-related preference for processing downward information, regardless of navigational relevance, in line with its underlying lower visual field bias.

The present study aimed at determining whether the OPA, PPA and MPA encoded information related to the vertical position of visual cues and whether this coding cared for navigational relevance. Specifically, we tested whether the vertical position of navigationally-relevant objects moderated the patterns of activity within scene-selective regions as a function of their retinotopic biases. We hypothesized that the OPA would show a preference for environments comprising cues in the lower visual field, regardless of their usefulness for navigation. We speculated that this effect in the OPA would be more pronounced in older adults than in young adults due to the downward processing bias that appears with age^18,45^. Moreover, we hypothesized that the PPA would exhibit a preference for environments encompassing cues in the upper visual field, irrespective of their usefulness for navigation. We anticipated that the MPA would not demonstrate any vertical preference. Accordingly, we designed a fMRI-based reorientation task that modulated the vertical position and task relevance of objects in a single-intersection virtual environment. We analyzed data using region-of-interest (ROI) univariate and multivariate pattern similarity analyses.

## Methods

### Participants

We enrolled 25 young and 26 older adults from the SilverSight cohort^49^ to partake in the current experiment. Participants in this cohort undergo a battery of neuropsychological, ophthalmological, auditory, and vestibular tests to ascertain that they are within the normal age-matched ranges for cognitive and sensory function. Moreover, they need to have normal or corrected-to-normal eyesight. A questionnaire requires individuals to indicate their sex assigned at birth and to confirm no history of neurological or psychiatric disorders. As part of the neuropsychological evaluation, participants complete the Mini-Mental State Examination (MMSE)^50^, the 3D mental rotation test^51^, and the perspective-taking task^52^. From the overall sample of 51 participants, we excluded 6 older adults due to poor task understanding and 1 young adult due to somnolence during the fMRI experiment. We therefore analyzed data from a group of 24 young participants (M = 28.08, SD = 4.03) comprising 13 women and 11 men, and 20 older participants (M = 73.95, SD = 5.12) comprising 10 women and 10 men. Enrolled individuals received monetary compensation in the form of a 30€ gift voucher. The Ethics Committee “CPP Ile de France V” reviewed and approved all experimental procedures (ID_RCB 2015-A01094-45, CPP N: 16122), and participants gave their written informed consent.

### Virtual navigation task

We designed the virtual environment with Unity3D v2019.2 (Unity Technologies, Inc. San Francisco, California, USA). It depicted a four-way intersection from which started four streets lined with identical buildings and sidewalks, making them strictly indistinguishable from each other. The first floor of each building comprised a balcony. At the level of the intersection, the sidewalks and balconies formed 8 right angles in total onto which objects could be positioned. We defined four conditions according to two factors: the number of objects and the vertical position of navigationally relevant objects. In *Half* conditions, the intersection contained a half set of objects located either on sidewalks (*Half* – DOWN) or on balconies (*Half* – UP). In *Full* conditions, the intersection contained a full set of objects with relevant objects located on sidewalks (*Full* – DOWN) or on balconies (*Full* – UP) and the 4 non-informative identical objects on balconies or sidewalks, respectively (Fig. 1). We used objects extracted from the Unity Asset Store based on their coherence with the virtual environment. We normalized the height of each item to 1 virtual meter and attenuated salient colors and textures. Prior to participant recruitment, 5 independent examiners assessed the recognizability of objects. We replaced objects judged to be poorly discernible by a majority of the raters. We created 32 unique versions of the intersection, 8 per condition, each with a new set of objects. We created an additional 32 instances that flipped the position of objects from sidewalks to balconies and from balconies to sidewalks. We conducted this procedure in order for participants to see each object in both vertical positions thereby attenuating possible variability linked to specific objects. In total, there were 64 unique instances of the environment.

**Figure 1.**
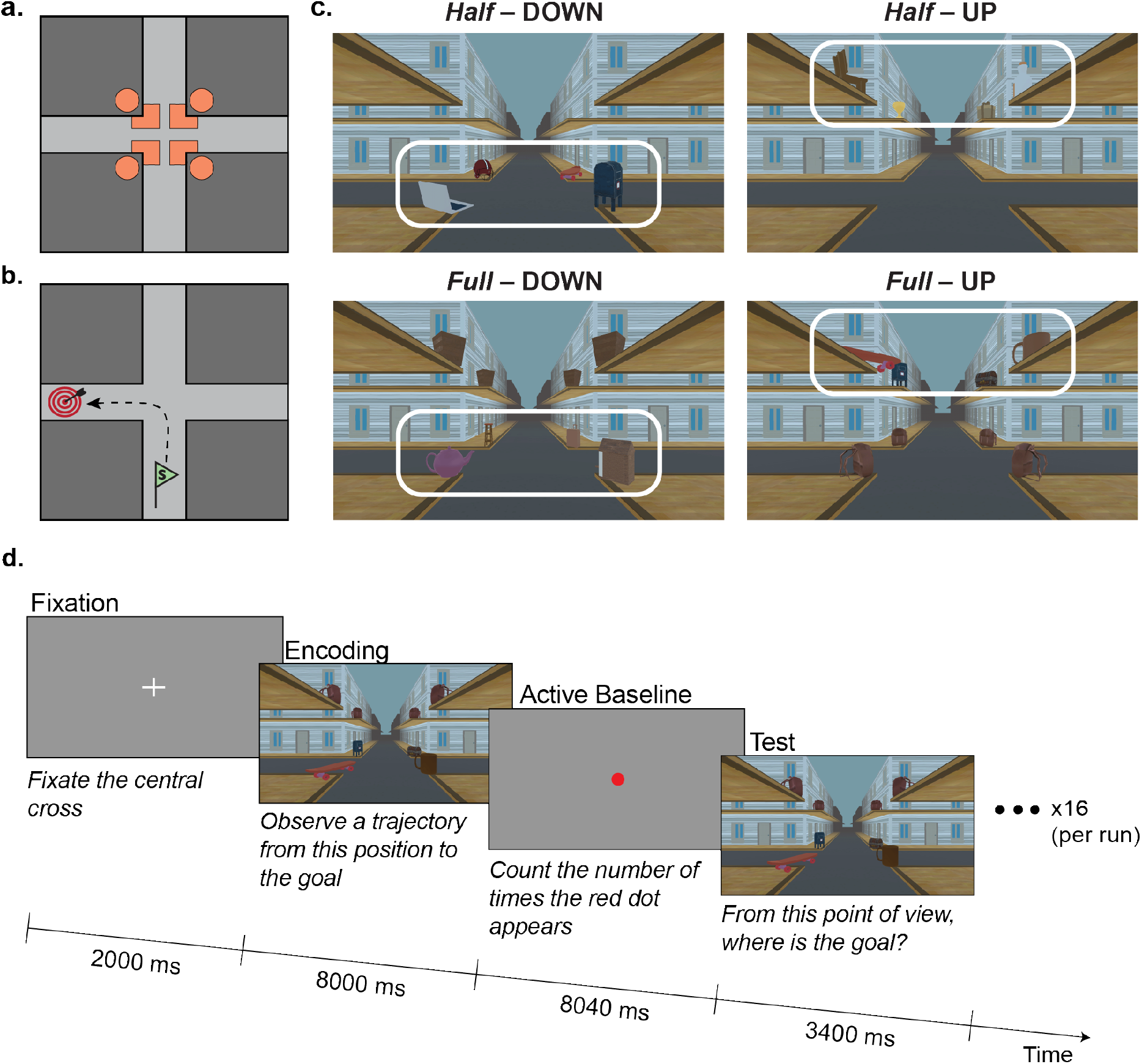
Description of the experimental paradigm. **a.** Overhead view of the 4-street intersection with the possible positions of objects indicated in orange. Circles represent the position of balcony objects and squares represent the position of sidewalk objects. **b.** Overhead view of the 4-street intersection with an example trajectory from the starting position (green flag) to the goal location (red target). **c.** Example intersections from the 4 conditions. The Half - DOWN condition contains objects on sidewalks only. The Half - UP condition contains objects on balconies only. The Full - DOWN condition contains objects that are useful for orientation on sidewalks and non-relevant objects on balconies. The Full - UP condition contains objects that are useful for orientation on balconies and non-relevant objects on sidewalks. The white boxes indicate the position of objects that are relevant for navigation. **d.** Schematic representation of a single trial within a run. There are 16 trials in a run. Participants first fixate a central cross. They observe a trajectory through an environment with a unique object configuration and they learn the position of the goal. Then, they count the number of times a red dot flashes during the active baseline task. Finally, they are put back into the previously learned environment in front of the intersection and they need to decide if the goal is located to their right or to their left.

Each trial was characterized by a unique version of the intersection and it consisted of 4 distinct phases. First, participants fixated a central white cross on a grey background for 2000 ms. Second, they performed the encoding phase of the navigation task. The latter projected participants into a first-person perspective and required them to observe a trajectory within the virtual environment. They started at the level of the intersection where they stayed for 3000 ms to allow sufficient time for item encoding. Then, the trajectory resumed and made a left or right turn into the street. The trajectory stopped upon reaching the goal (*i.e.,* a flower bouquet), 10 virtual meters into the street. Third, we included an active baseline task lasting approximately 8000 ms that asked participants to count the number of times a central black dot turned red. This task served as the baseline level of brain activity for comparison with activity during the navigation task. Finally, they performed the test phase of the navigation task. They were re-projected into a first-person perspective of the environment, facing the intersection either from the same learned direction or from the opposite direction. They had 3400 ms to decide whether the goal was situated in the street to their right or to their left by pressing a key. We did not provide participants with any feedback. Upon trial completion, a new trial started with the fixation screen followed by navigation in a different intersection. During the encoding and test phases of the navigation task, we set participants’ forward speed to 5 virtual meters per second and rotation speed to 67° per second.

### Functional localizer task

To precisely locate participants’ scene-selective regions, they underwent a functional localizer task, equivalent to that employed by previous studies^48,53^. It consisted of a block fMRI paradigm. We presented 900 x 900-pixel grayscale images subtending 18 x 18° of visual angle to participants for 400 ms. There was a 600-ms interval separating the presentation of each stimulus. We divided the experiment into 14 blocks lasting 20 s each. There were 4 blocks with photographs of objects, 4 blocks with scenes, 4 blocks with faces, 4 blocks scrambled objects and 4 blocks with a simple fixation cross. The order of blocks was pseudorandomized but it remained the same between participants. We asked participants to fixate the center of the screen throughout the task and to press a button on the response device when they perceived two consecutive photographs to be identical. Supplementary Table 1 displays each participant’s ROI coordinates.

### Procedure and apparatus

The experimental session began with a familiarization session outside of the MRI. The practice comprised 16 trials with objects not included in the main task. Participants repeated the practice until they reached a score of 70%. Upon completion, participants underwent a single MRI session. The latter comprised 8 runs of the navigation task with structural brain images taken halfway through and it ended with the functional localizer task. A single run consisted of 4 trials per condition (*i.e.,* 16 trials in total) displayed pseudorandomly with the rule that no consecutive trials could be from the same condition. In total, participants engaged in 128 trials corresponding to two possible starting positions within each of the 64 environment instances. We presented the original and flipped versions of the intersection as well as the two starting positions from the same environment in a single run. We ensured that half of the trials tested participants from the opposite point of view as that learned during the encoding phase. One run had an approximate duration of 6.2 mins. We counterbalanced the order of runs between participants using a balanced latin square design. Following the experimental session, we sent a Google Form questionnaire for participants to complete at home. It probed their use of objects present at the intersection and of the strategy they deployed to encode the goal position (Fig. S1).

We displayed the navigation and functional localizer tasks on a MRI-compatible liquid crystal display monitor (NordicNeuroLab, Bergen, Norway) that was positioned at the rear of the magnet bore. We presented stimuli from the navigation and functional localizer tasks and collected all responses in Nordic Aktiva v1.3.1 (NordicNeuroLab, Bergen, Norway) from a Dell Precision 3561 mobile workstation. Participants decided on the goal position using a MRI-compatible ergonomic two-grip response collection system (NordicNeuroLab, Bergen, Norway). They used left and right thumb presses to indicate that the goal position was to their left or to their right, respectively.

### Imaging parameters

We collected the imaging data at the Neuroimaging and Radiology Unit of the Quinze-Vingts National Hospital in Paris using a 3 Tesla Magnetom Prisma MR scanner (Siemens, Erlangen, Germany) equipped with a 64-channel head coil. For each run of the navigation task, we acquired 138 whole-brain images using T2*-weighted echo-planar imaging (EPI) sequences sensitive to the BOLD contrast with the following parameters: number of slices = 48; slice order = interleaved; repetition time (TR) = 2680 ms; echo time (TE) = 30 ms; voxel size: 3 x 3 x 2 mm^3^; matrix = 74 x 74; field of view (FoV) = 220 x 220 mm; flip angle = 90°. For the functional localizer task, we acquired 284 whole-brain images using T2*-weighted simultaneous multi-slice EPI sequences with the following parameters: number of slices = 64; slice order = interleaved; TR = 1000 ms; TE = 30 ms; voxel size: 2.6 x 2.6 x 2.4 mm^3^; matrix = 100 x 100; flip angle = 90° with a GRAPPA acceleration factor of 2. Finally, we acquired the structural three-dimensional brain image using a high-resolution T1-weighted Magnetization Prepared Rapid Gradient Echo (MPRAGE) sequence with the following parameters: number of slices = 176; TR = 2300 ms; TE = 2.98 ms; voxel size: 1 x 1 x 1.1 mm^3^; matrix = 256 x 248; flip angle = 9°. We oriented all slices along the long axis of the hippocampus.

### Behavioral analysis

We conducted the behavioral analyses using R version 4.0.3 in RStudio version 2022.12.0+353^54,55^. We used the *glmmTMB* package implemented in R to fit the generalized linear mixed models. The latter package uses maximum likelihood estimation and the Laplace approximation for random effects. We first studied the number of correct answers given by participants during the test phases to assess their navigational performance. We conducted a binomial generalized linear mixed model and performed model selection using the Akaike Information Criterion (AIC) goodness-of-fit statistic to identify the best fixed-effects and random-effects structure. The final model included condition, run number and age group as fixed effects along with participant and trial number as random intercepts. In accordance with guidelines pertaining to the modeling of reaction time data^56^, we performed a gamma generalized linear mixed model with a log link function to study the time it took participants to make a decision during the test phases. We detected and removed outliers from reaction time data using the interquartile range method. We built the model with condition, run number and age group as fixed effects and with participant and trial number as random intercepts. In addition, we used the post-experiment questionnaire to group participants into their preferred object position for orientation: sidewalks, balconies, or no preference. As an exploratory analysis, we performed a chi-squared test to verify whether the proportion of participants in each preference group differed between young and older adults.

### FMRI data preprocessing and analysis

We analyzed the imaging data using SPM12 (Wellcome Trust Centre for Neuroimaging, London UK, http://www.fil.ion.ucl.ac.uk/spm/) and custom codes in MATLAB R2022a. Before preprocessing, we discarded the first 4 scans from the localizer task to allow magnetization to reach steady state equilibrium. We also discarded the first 5 and last 5 scans from each run of the virtual navigation task. Of note, we focused the fMRI analyses on the encoding and not the test phase of trials as the former engages navigational processes and it is devoid of motor and decisional components.

#### ROI definition

We analyzed data from the functional localizer experiment in order to extract ROI masks for each participant. We first realigned all images to the first volume of the fMRI time series and we co-registered the functional images to the high-resolution anatomical T1-weighted scan. We then computed the forward deformation field from the T1 image to the Montreal Neurological Institute (MNI) space and used it to normalize the functional volumes. The following step consisted in resampling the images to a 3×3×3 mm voxel size by applying 4th degree B-spline interpolation. Finally, we smoothed the normalized volumes with an 8-mm full width half maximum Gaussian (FWHM) kernel. The preprocessing pipeline did not include slice timing correction as it is not recommended for multi-slice sequences^57^. To analyze the preprocessed localizer data, we used a general linear model (GLM) for block design with 5 regressors of interest convolved with the SPM canonical hemodynamic response function (HRF; *i.e.,* double gamma function). The 5 regressors corresponded to the 5 stimuli categories: scenes, faces, objects, scrambled objects and fixation. We also included the 6 rotation and translation movement parameters in the design matrix and we high-pass filtered the time-series from each voxel (1/128 Hz cut-off). We extracted the individual t-maps from the fMRI contrast [Scenes > Faces + Objects]. We then identified significant clusters, comprising at least 10 voxels, using family-wise error correction (ɑ = 0.05, t-value > 4.8). We extracted the masks for the left and right OPA, PPA, and MPA as the 40 contiguous voxels with the highest t-values around the peak activation. We then merged the ROIs from both hemispheres to create 80-voxel bilateral ROIs. Finally, we used the inverse deformation field to convert each ROI back into participants’ subject space and we resampled them to the voxel size of images from the navigation task (*i.e.,* 3×3×3 mm).

#### Univariate analyses

In a first analysis, we explored the fMRI data using ROI-based univariate analyses to gain insights into the average activation levels and specific differences between conditions. We first submitted the EPI images to slice-timing correction and within-subject realignment. Specifically, we realigned the functional scans to the first acquired BOLD image from that run to account for head motion. We then co-registered the images to the participant’s T1-weighted high-resolution structural image. Finally, we normalized the functional scans to the MNI space and we applied spatial smoothing with an 8-mm FWHM kernel. We implemented a motion scrubbing procedure using the CONN toolbox^58^. We computed the framewise displacement (FD) and we flagged volumes with FD > 0.9 mm, a common threshold in fMRI experiments with older populations^59,60^. We set the criterion that runs with less than 60% surviving volumes should be excluded from further analysis. As a consequence, we removed one run from an older participant. We fit the preprocessed data to subject-level GLMs with 18 regressors of interest for each run: (1) onsets and durations of the encoding phases from the 16 trials, (2) onsets and durations of the active baseline tasks from the 16 trials and (3) onsets and durations of the fixation phases from the 16 trials. We included the 3 rotation and 3 translation movement parameters as confound regressors. We also added a volume-wise regressor that modelled out volumes that were flagged during motion scrubbing. We convolved the GLMs with a double gamma function in order to best describe the shape of the HRF and we high-pass filtered (1/128 Hz cut-off) the time series from each voxel. In order to elucidate the patterns of activity within the OPA, PPA and MPA in relation to the experimental conditions, we extracted t-activation maps from the individual ROI masks for 6 different contrasts of interest. The main contrasts compared activity from the *Half* – UP, *Half* – DOWN, *Full* – UP, and *Full* – DOWN conditions to activity from the active baseline task. We also extracted activation differences between the *Half* – UP and *Full* – UP conditions and between the *Half* – DOWN and *Full* – DOWN conditions themselves. First, we implemented a linear mixed model to examine the effects of age group, ROI and condition on the t-maps associated with the 4 main contrasts. To further investigate the influence of the presence of non-informative objects, we conducted two separate linear mixed models with age group and ROI as fixed effects using the fMRI contrasts [*Half* – UP and *Full* – UP] and [*Half* – DOWN > *Full* – DOWN]. We included participants as a random intercept in all models.

#### Multivariate pattern similarity analyses

The main analysis of this study consisted in a multivariate pattern similarity analysis (MPS) using a Least-Squares Separate approach^61^. The latter enabled us to relate fine-grained patterns of activity across voxels to the cognitive processes subtending the encoding of information for navigation without interference from neighbouring task components. The preprocessing pipeline differed to that implemented for univariate analyses as it only included slice-timing correction, realignment and co-registration steps. We did not apply normalization or spatial smoothing to the functional images as it is not traditionally performed for single-trial estimations^62,63^. We estimated separate GLMs for each of the 128 trials with the encoding phase as a regressor of interest and all other encoding phases within the same run aggregated into a second regressor. We included the following regressors in the models: (1) encoding phase from the trial of interest, (2) encoding phases from all other trials within the same run, (3) test phase from the trial of interest, (4) fixation from the trial of interest. We also added the run-wise rotation and translation movement parameters as covariates of no interest. In order for the active baseline task to be considered as an implicit baseline, we left it unmodelled. Akin to univariate analyses, we convolved the GLMs with a double gamma canonical HRF. We extracted a t-map for every single estimated trial from the participants’ OPA, PPA and MPA masks and we assigned each t-map to its corresponding condition (i.e., *Half* – UP, *Half* – DOWN, *Full* – UP, *Full* – DOWN). We averaged the t-vectors across conditions, and we cast the mean values into a single matrix per ROI per participant. From these, we constructed representational similarity matrices (RSMs) by computing the Pearson correlation between pairs of conditions for each ROI and for each participant. In order to compare multivoxel patterns of neural activation between age groups we computed group-level RSMs by averaging the correlations through forward and backward Fisher z-transforms^64^. We therefore obtained 4×4 group-level matrices for the OPA, PPA and MPA that were symmetric around a diagonal of perfect correlations (*i.e.*, = 1).

We sought to investigate whether the neural RSMs reflected the absolute position of information or the position of relevant objects for navigation and if there were underlying preferences for a specific vertical location. To this end, we created 5 orthogonal RSMs that represented the expected similarity between conditions under various theoretical frameworks (Fig. S2). We constructed the *Absolute Position* RSM as a binary RSM by assigning 1 to condition pairs that had the same position of visual information at intersections (*i.e.*, *Full* – UP and *Full* – DOWN) and 0 to condition pairs that had differing object locations (i.e., *Half* – UP and *Half* – DOWN). The *Useful Position* RSM was also a binary RSM for which we assigned 1 to condition pairs that had useful objects located in the same vertical position (*i.e.*, *Half* – UP and *Full* – UP) and 0 to other pairs. We further constructed the *Upper Useful Position* and *Lower Useful Position* RSMs which we built upon the same principles as the *Useful Position* RSM with the additional constraint that they were specific for the upper or lower vertical positions respectively. Finally, we constructed the *Saliency* RSM in order to eliminate the possibility of imbalanced conditions in terms of object saliency. We calculated the objective saliency of each intersection screenshot using the graph-based visual saliency model implemented in Python (https://github.com/shreelock/gbvs) and we averaged the values across conditions. The final 4×4 *Saliency* RSM contained the Pearson correlation coefficients of average saliency values between each pair of conditions.

First, we ran a linear mixed model to study the influence of age group, ROI and pairwise comparison (*i.e.,* Does the correlation coefficient between *Half* – UP and *Half* – DOWN differ from that between *Full* – UP and *Full* – DOWN?) on the neural similarity patterns. We also conducted separate linear mixed effects models for each theoretical RSM to compare the neural RSM data against these prediction models. We fit the models in MATLAB R2022a using effects coding. Considering that the RSMs are symmetrical across the diagonal we only considered the lower triangular matrix in the models. We compared models with different random-effects and fixed-effects structures using the AIC and we started with a complex structure that still remained parsimonious^65^. This initial structure included Pearson correlation coefficients from the neural RSMs as the dependent variable. It comprised the theoretical RSM of interest, age group, and ROI along with their three-way interaction as fixed effects. We included ROI, pairwise comparisons between conditions and participants nested within age groups as random slopes. In the event of equivalent AIC values, we chose the best model based on the log likelihood. We performed two additional linear mixed models to pinpoint the theoretical RSM that best explained the neural RSM data in young and older adults separately. For this purpose, we computed the Pearson correlation coefficients between the RSM neural data in each ROI and the data from the 5 theoretical RSMs. These coefficients constituted the dependent variable in the models. We added theoretical RSM, ROI and their interaction as fixed effects, and we included participants as a random intercept.

To further characterize the distribution of neural patterns related to the vertical position of visual information, we conducted an exploratory whole-brain searchlight MPS. We built a sphere searchlight (6 mm radius, 33 voxels) centred on each voxel of the subject’s grey matter mask and we extracted the corresponding t-values from the spatial navigation task. We discarded spheres that were too close to the field of view boundaries or those that contained less than 20 voxels. Repeating the same analysis steps as described above, we constructed a RSM per searchlight sphere per participant. For each searchlight, we performed two-tailed permutation testing (ɑ = 0.05) on the voxel-level Fisher z-transformed Spearman rank correlation coefficients between the neural and theoretical RSMs. We thresholded the final group-level whole-brain maps at *p* < 0.001, smoothed them with a 6-mm FWHM Gaussian kernel and projected them onto a FreeSurfer inflated brain surface using Surf Ice v.6 (https://www.nitrc.org/projects/surfice/). Note that for the searchlight analysis we examined all theoretical RSMs except the *Saliency* RSM.

## Results

In the present study, 44 young and healthy older adults took part in a virtual navigation task while we recorded their neural activity with fMRI (Fig. 1). Participants observed trajectories in a first-person perspective through one-intersection environments. They needed to remember the position of a goal using objects positioned on the sidewalks and/or balconies of the intersection. Following an active baseline task, they were tested on their reorientation ability. The environments differed along two dimensions: the number of objects (*i.e.,* 4 or 8) and the vertical position of useful objects for orientation (*i.e.,* sidewalks or balconies). In *Half* – UP and *Half* – DOWN conditions, the intersection encompassed 4 objects located on balconies and sidewalks respectively. In *Full* – UP and *Full* – DOWN conditions, the intersection comprised 4 objects that were relevant for navigation on balconies and sidewalks respectively, and another 4 on the opposite vertical location that were not informative. We examined participants’ performance during the orientation task along with their subtending patterns of activity in the OPA, PPA and MPA. We performed both ROI-based univariate and multivariate pattern similarity analyses to compare the fMRI activity associated with parsing the vertical position of visual information for navigation in young and older adults.

### Behavioral results

We first ran a binomial generalized linear mixed model to examine how the proportion of correct responses per trial varied across conditions, age groups and runs. We found evidence for a main effect of age group on orientation accuracy (χ^2^(1) = 4.04, *p* = 0.045, *η*^2^ = 0.0055; Fig. 2a). Although this result indicates that older adults have significantly lower performance than young adults on the task, we need to highlight the very small effect size, in accordance with the relative simplicity of the task (M = 0.91 correct answers per trial, SD = 0.28 vs. M = 0.95, SD = 0.22). Importantly, there was no evidence for an influence of condition on behavioral performance in both age groups (χ^2^(3) = 2.22, *p* = 0.53, *η*^2^ = 0.00055), suggesting that participants can orient equally well across environments with varying object positions in the presence or absence of non-relevant navigational information (Fig. 2a). We also found a main effect of run number on mean accuracy (χ^2^(7) = 35.49, p < 0.001, *η*^2^ = 0.0064) reflecting a significant performance improvement between the first run and the following runs. For example, participants made on average 0.89 correct answers per trial during the first run and improved significantly to 0.96 during the third run (t(5442) = -4.86, *p* < 0.001, d = 0.13, SE = 0.053).

**Figure 2.**
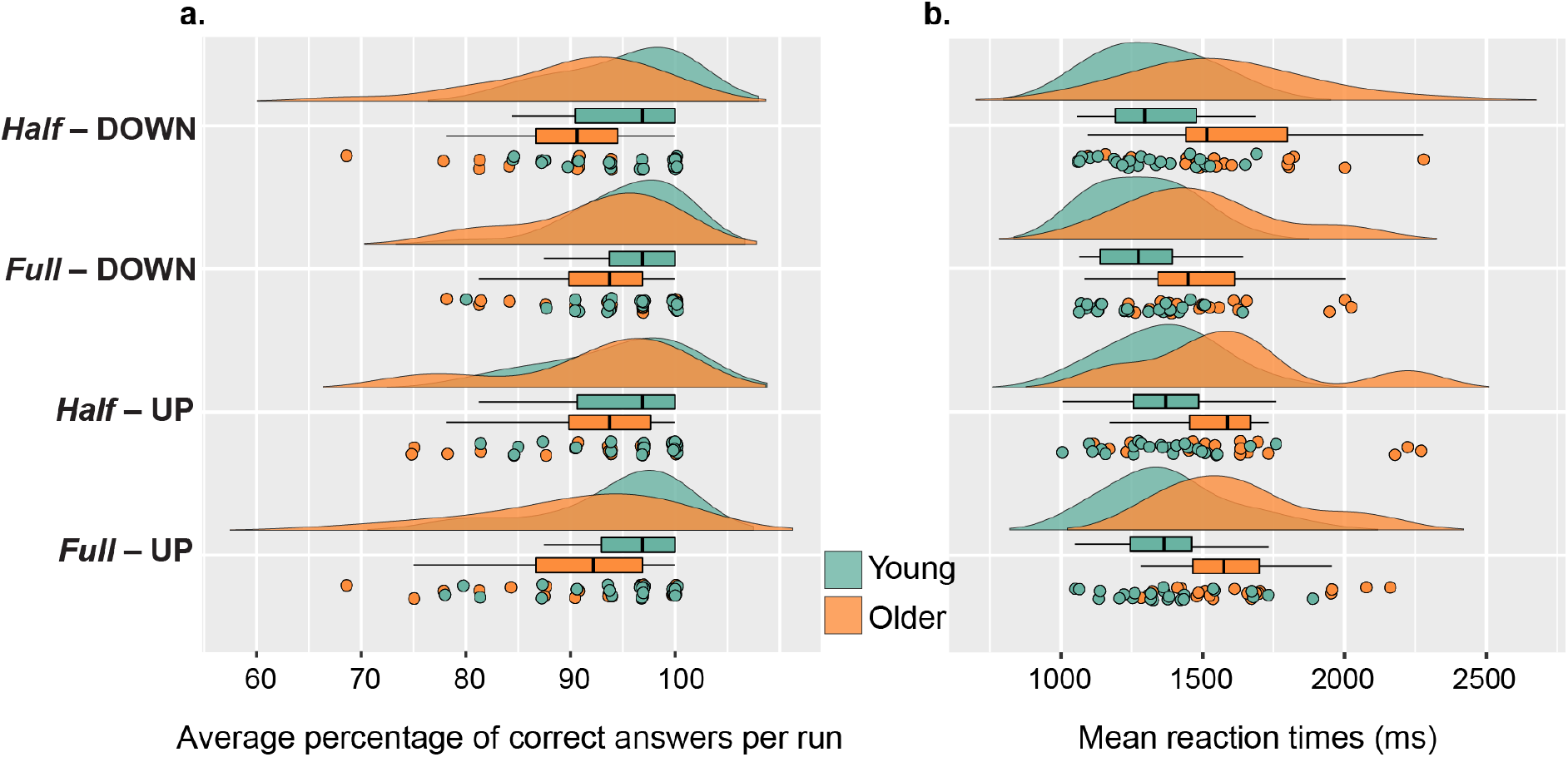
Behavioral performance during the test phases of the spatial navigation task. **a.** Distributions of the average percentage of correct answers per run across conditions in young and older adults. **b.** Distributions of the mean reaction times across all test phases in milliseconds (ms).

To further explore participants’ behavior, we ran a gamma generalized linear mixed model with a log link function to study the influence of condition, age group and run on reaction times during the test phases. First, we found a main effect of condition on reaction times (χ^2^(3) = 9.57, *p* = 0.023, *η*^2^ = 0.0076; Fig 2b). Post-hoc tests revealed that participants were slower to respond during *Full* – UP (M = 1469 ms, SD = 467 ms) than during *Full* – DOWN (M = 1360 ms, SD = 439 ms) conditions (t(4917) = -2.92, *p* = 0.018, d = 0.24, SE = 0.041). We also showed evidence for a significant effect of age group on reaction times (χ^2^(1) = 12.22, *p* < 0.001, *η*^2^ = 0.042; Fig. 2b). Older adults were slower to respond than young adults across all conditions (M = 1531 ms, SD = 483 ms *vs.* M = 1339 ms, SD = 432 ms). Finally, we found an influence of run number on reaction times (χ^2^(7) = 166.01, *p* < 0.001, *η*^2^ = 0.018) indicating that on average participants reduced the time they took to make a decision throughout the test phases of the experiment (Fig. S3).

### ROI-based univariate results

First, we wanted to compare the activity levels between the 4 different conditions to assess the interplay between the retinotopic biases of scene-selective regions and their responses to the position of navigationally-relevant objects during navigation. To this end, we performed a linear mixed model to explore the influence of condition, ROI and age group on the activity levels extracted from the fMRI contrasts [*Half* – UP > Active Baseline], [*Half* – DOWN > Active Baseline], [*Full* – UP > Active Baseline], [*Full* – DOWN > Active Baseline]. The univariate results revealed no evidence for a significant influence of condition on activity in scene-selective regions (F(3, 477) = 0.12, *p* = 0.95, *η_p_*^2^ = 0.00075, 95% CI [0.00, 0.0033]). This result underlines the idea that equivalent activation levels are required for spatial navigation in environments with and without task-irrelevant information. We showed however that ROI influenced activity levels (F(2, 477) = 56.16, *p* < 0.001, *η_p_*^2^ = 0.19, 95% CI [0.13, 0.25]; Fig. 3b). Post-hoc tests unveiled significant differences between the PPA and the OPA (t(477) = - 8.57, *p* < 0.001, d = 0.61, SE = 0.11) and between the PPA and the MPA (t(477) = 9.68, *p* < 0.001, d = 0.68, SE = 0.11). Indeed, there was greater activation in the PPA (M = 0.53, SD = 0.76) than in the OPA (M = 0.14, SD = 0.52) and MPA (M = 0.085, SD = 0.54) during the encoding phase of the navigation task across age groups and conditions. That increased PPA activity was found throughout conditions, regardless of cue position, is a first hint of an absence of retinotopic influence. Moreover, we found evidence for a main effect of age group (F(1, 42) = 14.57, *p* < 0.001, *η_p_*^2^ = 0.26, 95% CI [0.063, 0.45]; Fig. 3b). Overall, older participants (M = 0.50, SD = 0.59) displayed higher levels of activity than young participants (M = 0.043, SD = 0.62). Post-hoc tests revealed that these age-related differences stemmed from the OPA (t(61) = 3.07, *p* = 0.036, d = 0.83, SE = 0.16) and PPA (t(61) = 4.49, *p* < 0.001, d = 0.84, SE = 0.16), but not the MPA (t(61) = 2.87, *p* = 0.060, d = 0.74, SE = 0.16).

**Figure 3.**
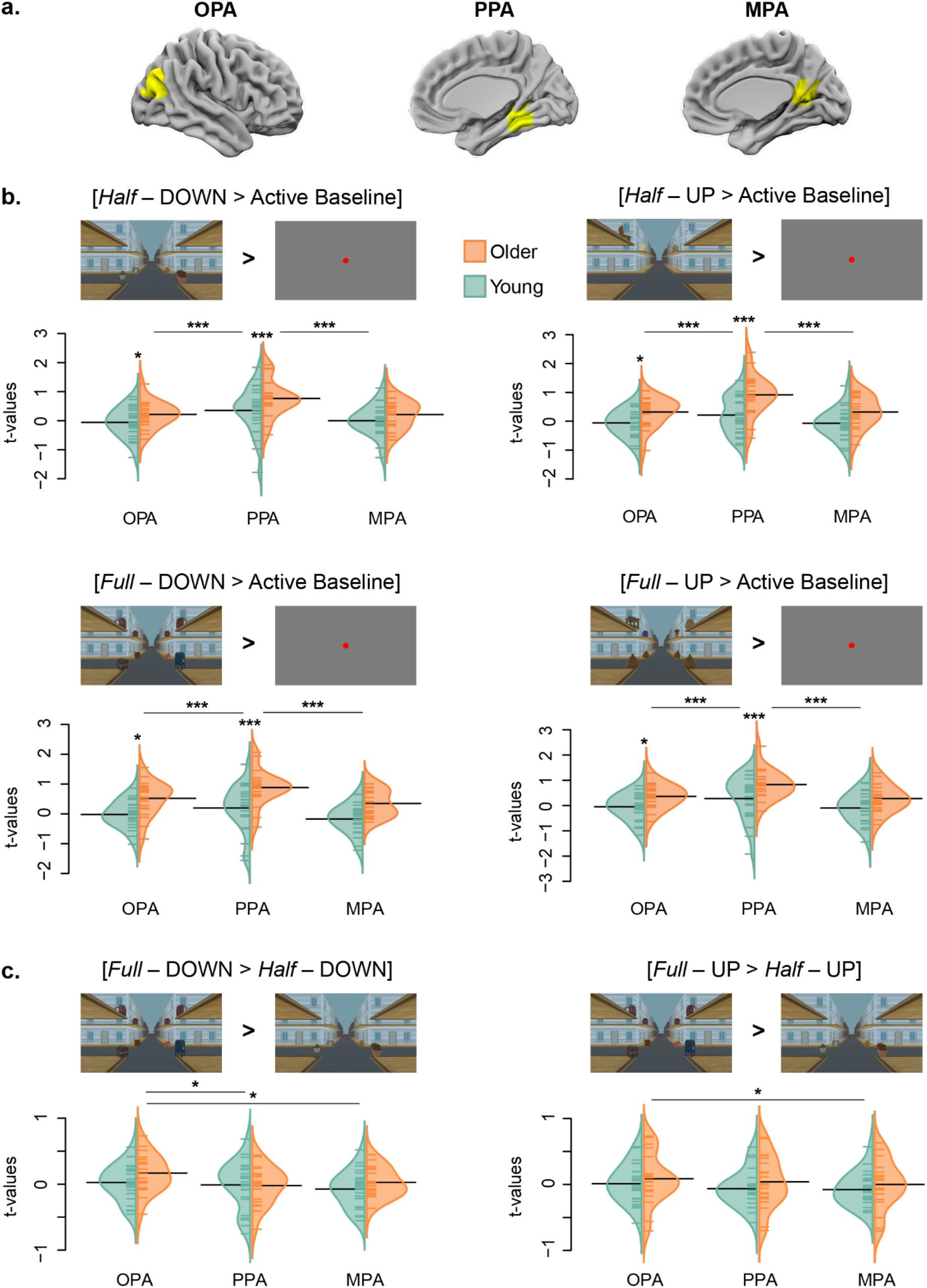
Results of the ROI-based univariate fMRI analyses. **a.** Illustrations of the localization of the OPA, PPA and MPA overlaid on the left hemisphere of one example participant. **b.** t-values extracted from the fMRI contrasts [*Half* – DOWN > Active Baseline], [*Half* – UP > Active Baseline], [*Full* – DOWN > Active Baseline] and [*Full* – UP > Active Baseline] in young and older participants. We conducted a linear mixed model to study the influence of condition, ROI and age group on fMRI activity levels. **c.** t-values extracted from the fMRI contrasts [*Full* – DOWN > *Half* – DOWN] and [*Full* – UP > *Half* – UP] in young and older participants. We ran two separate linear mixed models to study the influence of ROI and age group on fMRI activity differences between *Full* and *Half* conditions. *** p-value < 0.001, ** p-value < 0.01, * p-value < 0.05.

We next wanted to test the differences in activity between the *Full* and *Half* conditions for the UP and DOWN modalities. This analysis aimed to elucidate whether the absence of visual cues in a specific vertical hemifield influenced activation patterns within the OPA and PPA, in line with their respective retinotopic biases. Accordingly, we ran two linear mixed models to analyze the effects of ROI and age group on the neural activity extracted when contrasting *Full* – DOWN and *Half* – DOWN conditions and *Full* – UP and *Half* – UP conditions. First, we found a main effect of ROI on activity levels (F(2, 84) = 5.38, *p* = 0.0069, *η_p_*^2^ = 0.11, 95% CI [0.010, 0.24]; Fig. 3c), highlighting that neural activation in scene-selective regions is modulated by the use of lower objects in the absence or in the presence of non-informative upper objects. We further revealed with post-hoc tests that activity in the OPA was significantly different than that in the PPA (t(84) = 2.88, *p* = 0.014, d = 0.36, SE = 0.21) and MPA t(84) = 2.74, *p* = 0.020 d = 0.42, SE = 0.22). Contrasting with our initial hypothesis, the *Full* – DOWN condition elicited higher levels of activity than the *Half* – DOWN condition in the OPA (M = 0.090, SD = 0.28) but not in the PPA (M = -0.021, SD = 0.33) and MPA (M = -0.021, SD = 0.25). Of note, the difference between the OPA and PPA was primarily driven by the older adult group (t(84) = 3.31, *p* = 0.017, d = 0.70, SE = 0.33). We found no evidence for a main effect of age group on the activity levels extracted from the [*Half* – DOWN > *Full* – DOWN] contrast (F(1, 42) = 1.15, *p* = 0.29, *η_p_*^2^ = 0.027, 95% CI [0.00, 0.18]). Lastly, when contrasting the *Half* – UP and *Full* – UP conditions we also found a main effect of ROI on activity levels (F(2, 84) = 4.10, *p* = 0.020, *η_p_*^2^ = 0.089, 95% CI [0.0015, 0.21]; Fig. 3c), indicating that using upper objects in the absence or in the presence of non-informative lower objects elicited differential activations in scene-selective regions. Post-hoc testing revealed significant activity differences between the OPA and MPA only (t(84) = 2.79, *p* = 0.018, d = 0.27, SE = 0.21). It appeared that there was lower activity in the MPA (M = -0.042, SD = 0.31) than in the OPA (M = 0.045, SD = 0.33). We showed no evidence for a significant influence of age group on activation associated with the [*Half* – UP > *Full* – UP] contrast in scene-selective regions (F(1, 42) = 0.87, *p* = 0.36, *η_p_*^2^ = 0.020, 95% CI [0.00, 0.17]).

### ROI-based MPS results

The multivariate pattern similarity analysis strived to compare group-level neural RSMs in the OPA, PPA and MPA with orthogonal theoretical matrices. The main analysis consisted in deciphering whether the multivariate patterns of activation within the OPA, PPA and MPA reflected similarities between conditions sharing similar absolute positions of information or between conditions sharing similar useful positions of information. Furthermore, we also sought to investigate whether the observed patterns in these regions exhibited preferences for a specific vertical position in space. To this end, we conducted linear mixed models with theoretical model, age group and ROI as fixed effects to compare the neural similarity patterns against the 5 prediction RSMs separately (Fig. S2). We tested 2 theoretical models linked to processing incoming visual input (*i.e.*, *Absolute Position* and *Saliency* models) and 3 theoretical models linked to processing task-relevant position (i.e., *Useful Position*, *Upper Useful Position*, and *Lower Useful Position* models). Results pertaining to the *Absolute Position* and *Saliency* models suggest that the multivariate patterns of activation within the OPA, PPA and MPA don’t reflect incoming visual input. Further details can be found in Supplementary Note 1.

In order to test the hypothesis that scene-selective regions may rather be responsive to task relevance, we next strove to delineate the influence of the 3 theoretical RSMs related to processing the navigationally useful position of visual cues. We unveiled a main effect of the *Useful Position* prediction model (F(1, 786) = 55.50, *p* < 0.001, *η_p_*^2^ = 0.066, 95% CI [0.036, 0.10]) as well as an interaction between this theoretical RSM and ROI (F(2, 786) = 4.81, *p* = 0.0077, *η_p_*^2^ = 0.012, 95% CI [0.00086, 0.030]). Indeed, the *Useful Position* model explained the similarity patterns in the OPA to a greater extent than those in the PPA (F(1, 786) = 103.31, *p* < 0.001, *η_p_*^2^ = 0.12, 95% CI [0.078, 0.16]) and in the MPA (F(1, 786) = 170.89, *p* < 0.001, *η_p_*^2^ = 0.18, 95% CI [0.13, 0.23]). Regarding the *Upper Useful Position* model, we also observed a main effect of this theoretical RSM on neural similarity patterns (F(1, 786) = 19.77, *p* < 0.001, *η_p_*^2^ = 0.025, 95% CI [0.0077, 0.050]). Moreover, we found evidence for an effect of the interaction between the *Upper Useful Position* model and ROI (F(2, 786) = 6.51, *p* = 0.0016, *η_p_*^2^ = 0.016, 95% CI [0.0026, 0.037]). Similarly to the *Useful Position* model, the *Upper Useful Position* model accounted better for the similarity patterns in the OPA than in the PPA (F(1, 786) = 108.33, *p* < 0.001, *η_p_*^2^ = 0.12, 95% CI [0.082, 0.16]) and in the MPA (F(1, 786) = 161.97, *p* < 0.001). Finally, we unveiled a significant association between the *Lower Useful Position* model and the neural RSM data (F(1, 788) = 19.75, *p* < 0.001, *η_p_*^2^ = 0.025, 95% CI [0.0077, 0.050]). We found the latter association to be equivalent across ROIs. Overall, this analysis underlines the idea that scene-selective regions exhibit equivalent patterns of activation for environments that share the same position of task-relevant information.

To complement the previous analyses, we wished to delineate which theoretical model best explained the patterns of activation within the OPA, PPA and MPA of young and older participants. To this end, we ran a linear mixed model looking into the effects of theoretical model, ROI and the interaction between model and the ROI on correlation coefficients in young and older groups separately. In the younger group, we found evidence for a main effect of model (F(4, 322) = 17.20, *p* < 0.001, *η_p_*^2^ = 0.18, 95% CI [0.10, 0.24]; Fig. 4c). Post-hoc tests revealed significant differences between the *Absolute Position* model and the *Useful Position* (t(322) = -6.45, *p* < 0.001, d = 1.11, SE = 0.18), *Upper Useful Position* (t(322) = -4.87, *p* < 0.001, d = 0.79, SE = 0.17) and *Lower Useful Position* (t(322) = -5.45, *p* < 0.001, d = 0.88, SE = 0.17) models. Similarly, we found the correlation coefficients from the *Saliency* model to diverge from those from the *Useful Position* (t(322) = -5.74, *p* < 0.001, d = 0.91, SE = 0.18), *Upper Useful Position* (t(322) = -4.16, *p* < 0.001, d = 0.63, SE = 0.17) and *Lower Useful Position* (t(322) = -4.73, *p* < 0.001, d = 0.71, SE = 0.17) models. In young adults it was mostly the OPA that drove the differences between theoretical models (Table S1; Fig. 4c). In the older group, we also revealed that model had a significant effect on correlation strengths (F(4, 266) = 8.67, *p* < 0.001, *η_p_*^2^ = 0.12, 95% CI [0.045, 0.18]). Post-hoc tests revealed that there was a significant difference between the *Useful Position* model and the *Absolute Position model* (t(266) = 3.88, *p* = 0.0012, d = 0.69, SE = 0.19). We also found differences between the *Saliency* model and the *Useful Position* (t(266) = -5.21, *p* < 0.001, d = 1.00, SE = 0.19), *Upper Useful Position* (t(266) = -3.62, *p* = 0.0032, d = 0.66, SE = 0.19) and *Lower Useful Position* models (t(266) = -3.72, *p* = 0.0023, d = 0.71, SE = 0.19). Contrarily to young participants, in older participants it appeared that the MPA was primarily responsible for such differences (Table S2; Fig 4c). These findings confirm that in both young and older adults the activation patterns in scene-selective regions are best accounted for by equivalent encoding of environments that share the same position of useful information, irrespective of a specific vertical hemifield. Finally, we ran a complementary analysis to confirm that the neural RSMs differed between the three ROIs (Supplementary Note 2).

**Figure 4.**
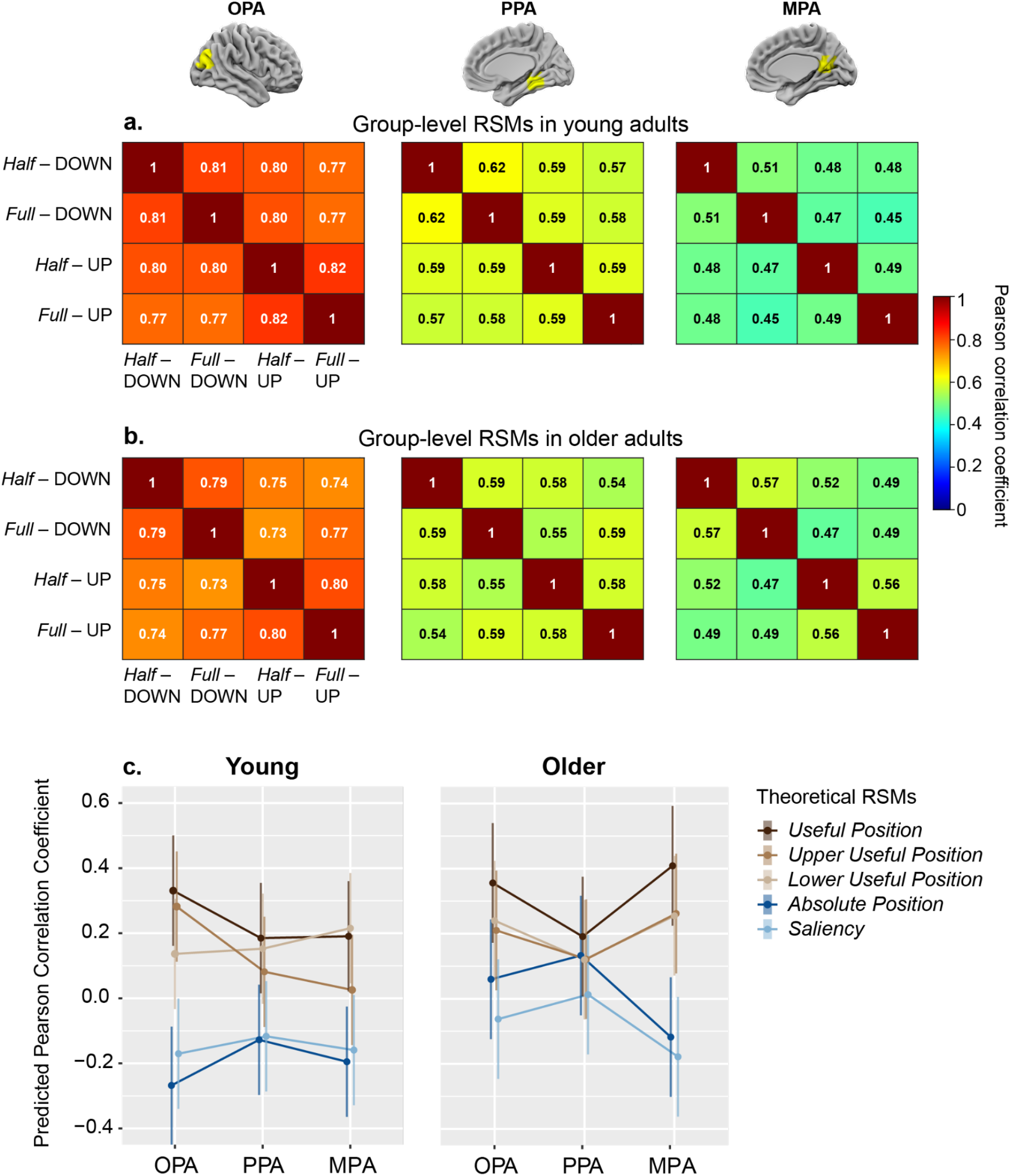
Results of the MPS analysis. **a.** Group-level representational similarity matrices (RSMs) in young adults for the OPA, PPA and MPA. Similarity scores are Pearson correlation coefficients, and they represent the overlap in neural patterns between the 4 conditions. **b.** Group-level RSMs in older adults for the OPA, PPA and MPA. **c.** Pearson correlation coefficients between the neural RSMs and 5 theoretical RSMs in the OPA, PPA and MPA. The *Useful Position*, *Upper Useful Position* and *Lower Useful Position* RSMs are linked to cognitive processing during the navigational task (in brown). The *Absolute Position* and *Saliency* RSMs are associated with the visual input at intersections (in blue).

### Searchlight-based MPS results

Finally, we conducted a searchlight analysis to explore whether this property of coding for the task-relevant position of visual cues extended to other brain areas. We did not find any cerebral region in young or older participants that overlapped with the *Absolute Position* RSM (Fig. 5a). However, we detected that the *Useful Position* RSM correlated significantly (*p* < 0.001) with activity in two main clusters of the occipital lobes, located on each side of the calcarine fissure, in both age groups (Fig. 5b). In young and older participants, we found the peak voxels from the dorsal clusters to be located in the left (young: [x = -6, y = -91, z = 24], t = 8.34; older: [-6, -87, 21], t = 7.23) and right cuneus (young: [6, -84, 23], t = 7.10; older: [12, -89, 25], t = 7.54). The peak voxels from the ventral clusters on the other hand were situated along the calcarine fissure in older adults ([-1, -87, 6], t = 8.58) and deeper in the left ([-10, -79, -9], t = 5.40) and right lingual gyri ([9, -72, -3], t = 5.40) in young adults. We identified a supplementary cluster of activity in older adults that was absent in younger adults, located higher along the visual dorsal stream in the left precuneus ([-6, -66, 54], t = 3.87). We found the *Upper Useful Position* RSM to overlap with clusters along the dorsal stream only (Fig. 5c). We detected the peak voxels from the main clusters to be located in the left (young: [-6, -93, 21], t = 7.15; older: [-6, -90, 25], t = 9.11) and right cuneus (young: [12, -87, 35], t = 8.28; older: [7, -90, 19], t = 10.21) for both age groups. The *Upper Useful Position* RSM also correlated significantly with a more anterior cluster of the dorsal stream along the parieto-occipital fissure in the young group ([0, -81, 42], t = 7.57) and in the left precuneus in the older group ([-3, -69, 59], t = 5.35). Finally, we identified clusters in the ventral stream whose patterns of activity overlapped with the *Lower Useful Position* RSM (Fig. 5d). In young adults we found the peak voxel to be on the calcarine fissure ([0, -84, -6], t = 6.46). In older adults we identified the peak voxels to be positioned more anteriorly in the left ([-9, -66, 3], t = 4.99) and right lingual gyri ([6, -63, 3], t = 5.11).

**Figure 5.**
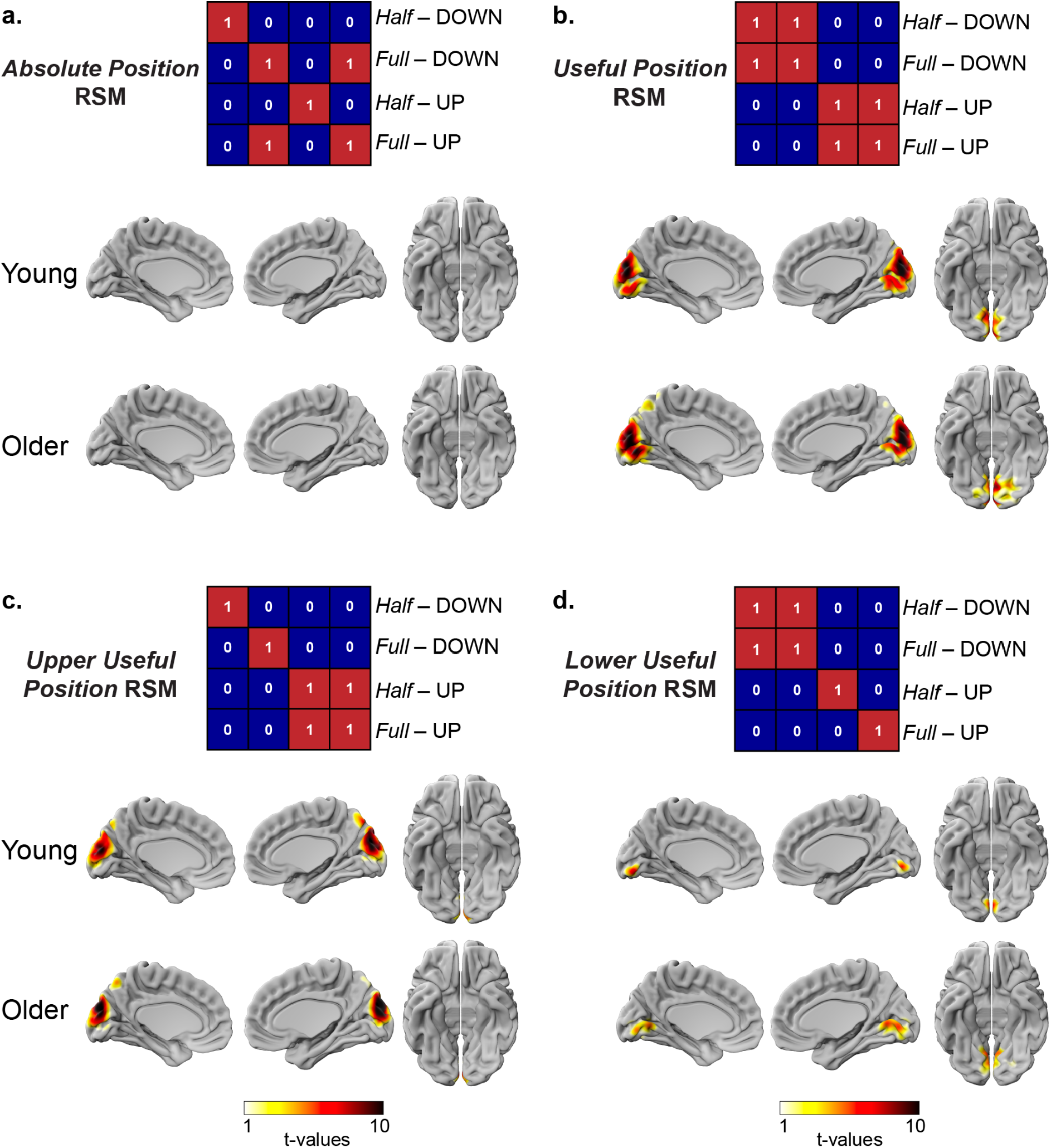
Results of the searchlight MPS analysis. Left hemisphere, right hemisphere and bilateral inflated surfaces showing the t-value of the searchlight centre voxel from the regions whose patterns of activity correlated with the **a.** *Absolute Position* theoretical representational similarity matrix (RSM), **b.** *Useful Position* RSM, **c.** *Upper Useful Position* RSM and **d.** *Lower Useful Position* RSM. t-maps were thresholded at *p* < 0.001.

## Discussion

The present study aimed at delineating the significance of the vertical position of task-relevant visual cues, and its interaction with the retinotopic biases of scene-selective regions during a spatial navigation task. We also examined whether patterns of activation in response to specific vertical locations varied between young and healthy older adults. Specific attention was directed towards the documented preference of older individuals for the lower visual field. First, we revealed that objects in the lower part of a scene may facilitate performance by lowering reaction times. We then showed that the OPA, and to a lesser extent the PPA and MPA, parsed the vertical position of useful information for navigation in both age groups, irrespective of the underlying retinotopic biases. Despite this common coding framework, there were striking differences between the scene-selective regions within and across age groups, highlighting their distinct functions. This work also shed light on a clear demarcation between the ventral and dorsal visual streams regarding the processing of upper and lower task-relevant visual cues.

### Responsiveness of scene-selective regions to the position of task-relevant information

Expanding upon the behavioral findings revealing a performance advantage when navigational information was situated in the lower part of the scene during *Full* conditions, the fMRI results put forward the dual coding of vertical position and task relevance of objects for effective spatial navigation. Indeed, the MPS analysis revealed that scene-selective regions extract the vertical position of task-relevant information in a navigable space in young and older adults. In other words, they produced similar patterns of activity when navigationally relevant objects were located in the same position, in contradiction with our initial hypothesis. We expected that scene-selective regions would demonstrate sensitivity to objects at positions in space corresponding to their retinotopic biases, regardless of task relevance. This study nonetheless aligns with two parallel lines of research pertaining to scene-selective regions. The first concerns the critical role of scene-selective regions in processing navigation-related information^3,31,32,38,66–70^. For example, it was shown that scenes with comparable structural arrangement of paths evoked similar multivoxel activation patterns in the OPA^66^. The second line of research relates to a fundamental attribute of the scene-selective network, namely its subtending representation of 2D spatial location. While most studies document that the OPA, PPA and MPA carry representations of the retinotopic position of information^28,29,71,72^, others advocate for robust representations of the perceived position of information^73^. As the current experiment was conducted under ecological free viewing conditions, we argue that scene-selective regions are indeed capable of encoding the position of information in spatiotopic coordinates. This hypothesis gains further support from our finding that the lower retinotopic bias in the OPA and the upper retinotopic bias in the PPA did not manifest as preferences for either task-relevant or task-irrelevant object positions in the context of navigation. The OPA, PPA and MPA may assign greater significance to the position of stimuli when they hold navigational value, irrespective of the subtending retinotopic biases. Our study acts as a bridge between the two lines of research by revealing the blending of information related to navigation and to position within scene-selective regions.

### Vertical position of landmarks as a fundamental coding property of the visual system

The present findings emphasize the importance of the vertical dimension as a coding property of the visual system. Not only did we show that the scene-selective areas extract vertical position information from objects that hold navigational relevance but also that the ventral and dorsal streams process navigationally relevant information depending on its vertical spatial location. The searchlight analysis revealed a strict dichotomy between the ventral and dorsal streams, with the former processing useful objects situated in the lower portion of the screen and the latter processing useful objects in the upper portion of the screen. These results resonate with previous work unveiling similar neural representations in the visual system for parts of scenes that occupy identical locations along the vertical axis^74,75^. We add further compelling empirical support to the hypothesis formulated by Kaiser and Cichy (2021)^76^ that the human brain is wired to sort visual inputs according to their upper or lower positions. An interesting question resides in understanding why verticality is an important cue for scene processing and spatial navigation. One possibility is that the summary statistics in the vertical dimension provide more informative cues about a scene compared to those in the horizontal dimension. For example, a recent study established the vertical luminance gradient to be a reliable discriminatory feature of a scene as light falls differently in the upper and lower visual fields^77^. Another non-exclusive explanation pertains to the navigational relevance of cues typically found in the vertical hemifields. A previous investigation looking at the sensitivity of scene-selective areas to position-in-depth unraveled a bimodal preference for front and back depths^78^, agreeing with the bimodal coding of upper and lower useful positions observed in the current study. Here we demonstrate that in the absence of depth differences between the upper and lower navigational cues, the visual system still encodes the two vertical positions separately. The vertical location of stimuli could therefore constitute a cue in itself. The strict division of activation elicited by upper and lower navigational cues in the dorsal and ventral visual streams, respectively, gives rise to compelling interrogations. First, this finding cannot be explained by underlying retinotopic maps as the dorsal stream receives more information from the lower visual field whereas the ventral stream predominantly receives information from the upper visual field^25^. We observed that processing navigationally relevant objects located in the upper portion of the screen required a widespread pattern of activation that reached anterior regions of the dorsal stream. The fMRI response accompanying the processing of task relevant objects from the lower portion of the screen was considerably more localized, with peak activity in the lingual gyrus. Such distinct patterns echo with recent literature putting forward the idea that object categorization recruits the dorsal stream when stimuli are unconventional or challenging^79^. In line with the behavioral results showing longer reaction times during *Full* – UP conditions, it may the case that the atypicality of a scene requires more intensive neural computations along the dorsal stream. Finally, we wish to stress that our paradigm, much like previous experiments examining near and far distances^80^, was designed such that only two vertical positions could be tested. As such, additional investigations are essential to elucidate whether the vertical position of task-relevant information is encoded in a bimodal fashion or follows a continuous gradient.

### Insights into the encoding of vertical position in older adulthood

First, despite uncovering an age-related decline in wayfinding accuracy and reaction times across the various conditions, the magnitude of the impairment was very small. Univariate analyses showed that older adults still displayed higher activation levels in the OPA and PPA during the encoding phase across all 4 conditions compared to young participants. Our results align with those from a previous report that demonstrated comparable performance levels between age groups and an age-related increase in OPA activity during active navigation^48^. Accordingly, we also uncovered no evidence for age-related differences in neural similarity patterns suggesting that the specific roles of the OPA, PPA and MPA are maintained in older age. In light of these observations, we hypothesize that the activity of scene-selective regions is subject to modulation by older adults with the objective to attain normal performance levels on a given task. Moreover, the searchlight analysis revealed that older adults, much like young adults, processed upper and lower navigationally relevant information separately in the dorsal and ventral visual streams. The patterns of activation showed striking similarities between age groups, with the exception of one cluster identified for the *Lower Useful* RSM. In response to useful objects in the lower part of the scene, young participants displayed a cluster of activity within the posterior portion of the calcarine fissure. This region is known to be implicated in the processing of high-resolution foveal information^81^. In contrast, older participants exhibited a cluster that was shifted anteriorly, aligning more closely with regions associated with representations of peripheral space. This pattern of results echoes with previous research indicating impaired fine-grained processing and hypoactivation of cerebral regions associated with foveal processing in older individuals^82,83^.

We argue that the preservation of scene-selective regions throughout adulthood strengthens the case for the importance of segmenting space into upper and lower vertical dimensions in scene processing and spatial navigation^22^. We stress that despite mounting evidence for a lower visual field bias in healthy older adults, neural representations across the visual system were equivalent for objects in the upper or lower part of the visual field^18,43–45^. Finally, the most noteworthy age-related difference that we reported pertains to the correlation coefficients between neural and theoretical RSMs. Specifically, we found that correlation coefficients were less distinct between the five theoretical models and the neural RSM in older participants than in young participants. This finding aligns with past literature documenting neural dedifferentiation in scene-selective regions in healthy aging^84,85^. Notwithstanding a relative maintenance of neural representations within the OPA, PPA and MPA of older individuals, it is conceivable that the distinctiveness of patterns of activation is reduced.

### Limitations and perspectives

Despite compelling evidence supporting verticality as an essential cue for spatial navigation, we must acknowledge the potential influence of horizontal position on both behavior and neural patterns throughout adulthood. It would be highly pertinent for future studies to explore the effect that varying positions along the horizontal meridian could have on the activity of scene-selective regions. The results obtained with the searchlight analysis raise an intriguing question: could the ventral and dorsal streams also segregate useful objects according to their position along the horizontal dimension? While we carefully designed the task to mitigate behavioral disparities between young and older adults, it is important to acknowledge that the simplicity of the paradigm may be viewed as a limitation in certain respects. Indeed, a more complex spatial navigation task is warranted to unearth potential behavior and neural differences linked to the use of visual cues in the upper or lower visual field. Finally, it would have been valuable to include eye tracking measures in this experiment in order to elucidate the relationship between the fine-grained attentional profile of participants and their fMRI patterns of activation.

### Conclusions

In summary, the present study unveils a role for the scene-selective regions, and particularly the OPA, in linking the vertical location of information with task relevance during a spatial navigation task. This property of scene-selective regions to encode vertical position applies solely to task-relevant objects and is independent of subtending retinotopic biases. Moreover, we revealed that the processing of navigationally relevant objects in the dorsal and ventral visual streams is strictly contingent upon their vertical location in space. Task-relevant objects in the upper part of the visual field are processed dorsally whereas task-relevant objects in the lower part of the visual field are processed ventrally. We further found that the influence of verticality on neural computations in the visual system remains intact in older age. We argue that the position of navigationally relevant information along the vertical dimension constitutes a fundamental coding property of the visual system throughout adulthood that may facilitate landmark-based spatial navigation. Overall, this study sheds new light on the intricate relationship between vision and spatial navigation and it highlights the importance of considering the position of stimuli for adequate cognitive functioning in real-world environments.

## Data availability

The datasets generated and analyzed during the current study are available via the OSF repository: https://osf.io/5m3p7/?view_only=21107e06a58e47ac9220305769c67f28.

## Code availability

Custom codes used to run the behavioral and neuroimaging analyses and to plot the results are available at: https://osf.io/5m3p7/?view_only=21107e06a58e47ac9220305769c67f28. We wrote the codes using R version 4.0.3 in RStudio version 1.4.1103 (R Core Team, 2020; RStudio Team, 2021) and Matlab R2022a (Mathworks Inc., Natick, MA, USA).

## Acknowledgements

We are deeply grateful to all the participants involved in this experiment. We thank Fabienne Tzvetkov-Ricard and Sonia Combariza from the Aging in Vision and Action laboratory at the Vision Institute for their assistance in enrolling participants. This research was supported by the French National Research Agency (ANR-18-CHIN-0002), the LabEx LIFESENSE (ANR-10-LABX-65), the IHU FOReSIGHT (ANR-18-IAHU-01), and the Fondation pour la Recherche sur Alzheimer (FRA).

## Author contributions

M.D., E.S., A.D., A.A. and S.R. designed the experiment. M.D., E.S. and L.L. collected the data. M.D. and L.L. analyzed the data. M.D. wrote the original manuscript. M.D., L.L., A.A. and S.R. revised the manuscript.

