## Supplementary Information for "Scene-selective regions encode the vertical position of navigationally relevant information in young and older adulthood"

**Supplementary Note 1**

We concentrated on testing the group-level RSMs against the theoretical RSMs associated with visual input. We first found that there was little evidence to suggest an effect for the *Absolute Position* RSM ($F(1, 786) = 0.00024$, $p = 0.99$, $\eta_p^2 = 0.00$, 95% CI [0.00, 0.00]). However, we revealed a significant influence of the interaction between this theoretical model and age group ($F(1, 786) = 9.64$, $p = 0.0020$, $\eta_p^2 = 0.012$, 95% CI [0.0016, 0.032]). While the effect size is small, this result underlines the fact that the *Absolute Position* RSM was differentially associated with the neural RSMs in older and young participants. We then investigated whether the saliency of objects at intersections could contribute to the neural similarity patterns. We found evidence that the *Saliency* theoretical model had an effect on the neural RSMs ($F(1, 788) = 11.03$, $p < 0.001$, $\eta_p^2 = 0.014$, 95% CI [0.0023, 0.034]). It is key to underline that the coefficient estimate for this effect was equal to -0.11 suggesting that when the neural similarity between conditions increased, the saliency similarity decreased. Thus, the *Saliency* model had no meaningful influence on the neural RSM data.

**Supplementary Note 2**

We ran a linear mixed model to decipher the effects of ROI, age group, and pairwise comparison between conditions on the multivariate patterns of activation. We found evidence for a main effect of ROI ($F(2, 778) = 94.60$, $p < 0.001$, $\eta_p^2 = 0.20$, 95% CI [0.15, 0.24]; Fig. 4). The similarity with which the 4 conditions were encoded differed between the OPA and PPA ($F(1, 778) = 93.28$, $p < 0.001$), OPA and MPA ($F(1, 778) = 180.10$, $p < 0.001$, $\eta_p^2 = 0.19$, 95% CI [0.14, 0.24]), and PPA and MPA ($F(1, 778) = 10.11$, $p = 0.0015$, $\eta_p^2 = 0.013$, 95% CI [0.0019, 0.033]). This result emphasizes considerable disparities in the correlation strengths between conditions in each of the 3 scene-selective regions. Notably, there was little evidence for an effect of age group on neural RSMs ($F(1, 778) = 0.027$, $p = 0.87$, $\eta_p^2 = 0.00$, 95% CI [0.00, 0.00]; Fig. 4). Overall, the similarity patterns appeared to be equivalent between young and healthy older participants across ROIs.

We examined how specific pairwise comparisons (e.g., *Half* - DOWN x *Full* - DOWN *vs. Half* - UP x *Full* - UP) modulated the correlation strength. The results are presented in Table S3. We revealed that pairwise comparisons had an influence on the neural RSMs ($F(5, 778) = 13.06$, $p < 0.001$, $\eta_p^2 = 0.077$, 95% CI [0.041, 0.11]; Fig. 4.2.4a,b). The correlation coefficients between *Full* – DOWN and *Full* – UP conditions did not differ significantly from the coefficients between *Half* – DOWN and *Half* – UP (F(1, 778) = 1.27, $p$ = 0.26, $\eta_p^2$ = 0.0016, 95% CI [0.00, 0.012]). Similarly, we found no evidence that correlation coefficients between *Half* – DOWN and *Full* –DOWN conditions differed from those between *Half* – UP and *Full* – UP conditions (F(1, 778) = 0.16, $p$ = 0.69, $\eta_p^2$ = 0.00021, 95% CI [0.00, 0.0068]). We revealed all other pairwise comparisons to be statistically dissimilar.

### Supplementary Figures

#### Questionnaire Post-Expérience

##### Difficulté de l'expérience

**1 ) Comment avez-vous trouvé cette expérience ?**

Très facile ○ 1 ○ 2 ○ 3 ○ 4 ○ 5 Très difficile

**2 ) Comment avez-vous trouvé l'utilisation des manettes ?**

Très facile ○ 1 ○ 2 ○ 3 ○ 4 ○ 5 Très difficile

**3 ) Les objets présents aux intersections vous ont-ils aidés à vous orienter ?**

☐ Oui

☐ Non

##### Objets et orientation

**4a ) Si oui, comment choisissiez-vous les objets qui vous aidaient à vous orienter ?**

☐ Je choisissais ceux que je préférais

☐ Je choisissais ceux qui se situaient le plus proche de moi

☐ Je choisissais ceux qui se situaient le plus loin de moi

☐ Je choisissais ceux qui étaient le plus visible pour moi

☐ Je choisissais toujours ceux de gauche

☐ Je choisissais toujours ceux de droite

☐ Je les choisissais tous (je retenais les 4 objets présents)

**4b ) Si non, comment faisiez-vous pour retrouver le bouquet de fleurs ?**

.......................................................................................................................

##### Environnements avec quatre objets

*Les questions suivantes ne concernent que les environnements où 4 objets étaient présents (soit 4 sur les balcons, soit 4 sur les trottoirs)*

**5 ) Avez-vous remarqué que les objets se situaient parfois sur les trottoirs et parfois sur les balcons ?**

☐ Oui

☐ Non

**6 ) Si oui, avez-vous trouvé un type d'environnement plus difficile qu'un autre pour retrouver le bouquet ?**

☐ Oui, j'ai eu plus de difficultés lorsque les objets étaient sur les balcons

☐ Oui, j'ai eu plus de difficultés lorsque les objets étaient sur les trottoirs

☐ Non

##### Environnements avec huit objets

*Les questions suivantes ne concernent que les environnements où 8 objets étaient présents (4 sur les trottoirs et 4 sur les trottoirs)*

**7 ) Selon vous, les 8 objets étaient-ils tous utiles pour retrouver le bouquet de fleurs ?**

☐ Oui

☐ Non

**8 ) Si non, selon vous, pourquoi certains objets ne pouvaient pas vous aider à retrouver le bouquet ?**

.......................................................................................................................

##### Questions supplémentaires

*Prenez en compte tous les différents types d'environnements (avec 4 ou 8 objets)*

**9 ) Avez-vous eu une préférence pour un environnement en particulier ?**

☐ Oui

☐ Non

**10 ) Décrivez la stratégie que vous auriez adopté face à ces intersection (le bouquet de fleurs se trouve à droite sur la première image et à gauche sur la deuxième)**

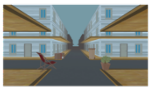
.......................................................................................................................

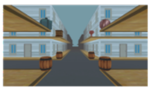
.......................................................................................................................

**Figure S1.** Questionnaire (in French) sent to participants as a Google form after the experiment. It probed their object preferences and strategy use for orientation in the virtual environment.

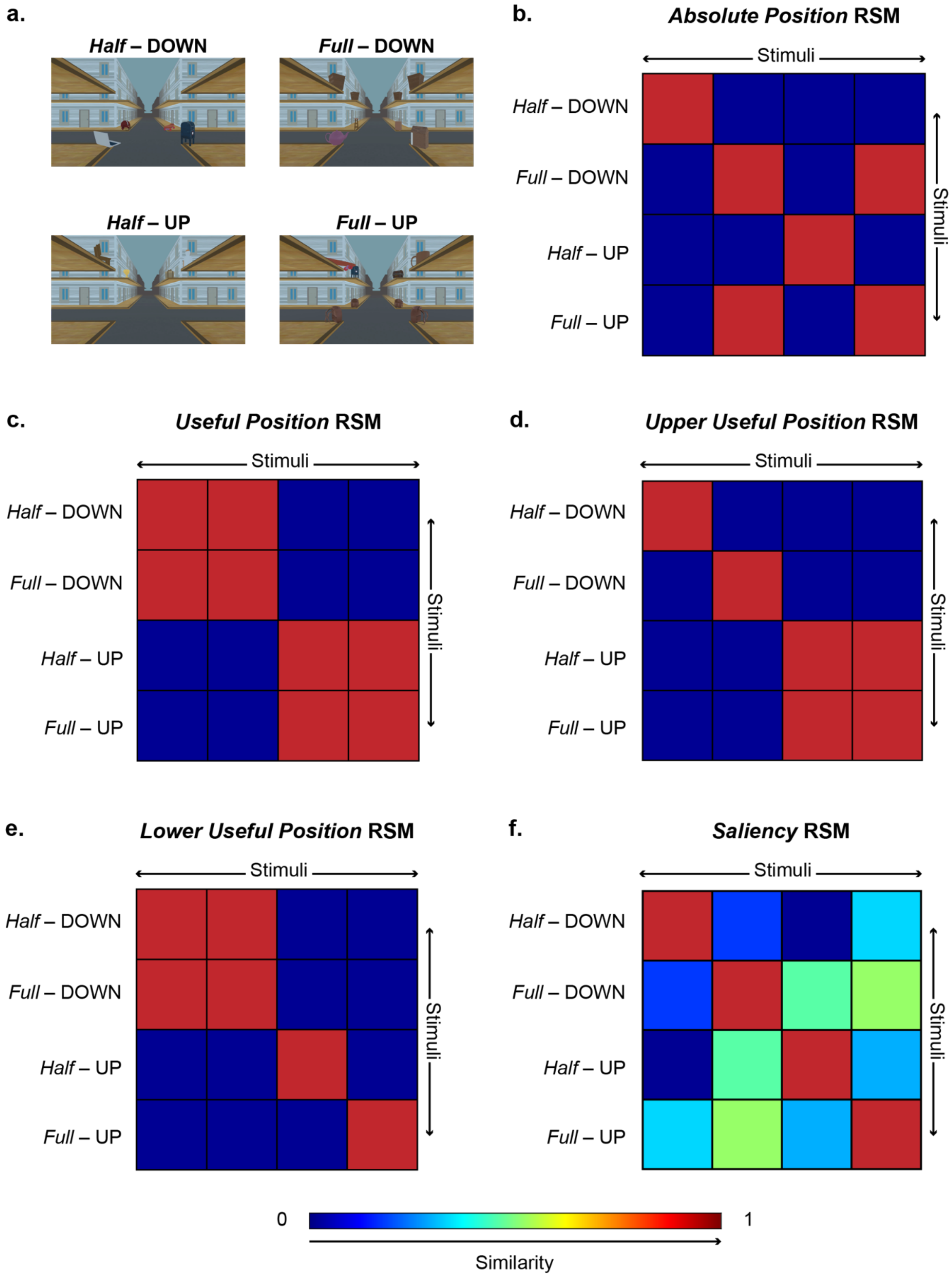


**Figure S2.** Orthogonal theoretical RSMs. **a.** Examples of intersections from the 4 conditions. **b.** The *Absolute Position* RSM considers conditions to be highly similar if objects at the intersection are in the exact same position. **c.** The *Useful Position* RSM considers conditions to be highly similar if objects that are relevant for orientation are in the same vertical position. **d.** The *Upper Useful Position* RSM considers conditions to be highly similar if objects that are relevant for orientation are on balconies. **e.** The *Lower Useful Position* RSM considers conditions to be highly similar if objects that are relevant for orientation are on sidewalks. **f.** The *Saliency* RSM considers conditions to be highly similar if the graph-based visual saliency values of their intersections are correlated.

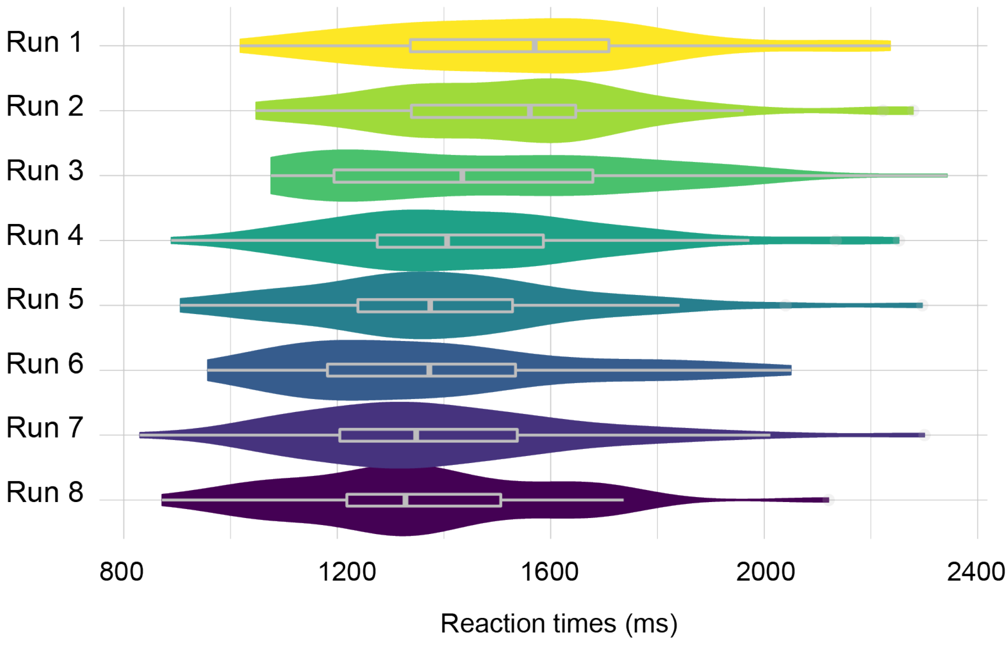


**Figure S3.** Reaction times across the 8 runs of the experimental paradigm. The reaction times in ms are averaged across young and older participants.

#### Supplementary tables

| | OPA | PPA | MPA |
|---|---|---|---|
| *Absolute Position > Useful Position* | t(322) = -5.16<br>$p$ < 0.001 | t(322) = -2.69<br>$p$ = 0.31 | t(322) = -3.32<br>$p$ = 0.067 |
| *Absolute Position > Upper Useful Position* | t(322) = -4.74<br>$p$ < 0.001 | t(322) = -1.80<br>$p$ = 0.90 | t(322) = -1.90<br>$p$ = 0.85 |
| *Absolute Position > Lower Useful Position* | t(322) = -3.49<br>$p$ = 0.041 | t(322) = -2.41<br>$p$ = 0.51 | t(322) = -3.54<br>$p$ = 0.034 |
| *Absolute Position > Saliency* | t(322) = -0.84<br>$p$ = 1.00 | t(322) = -0.093<br>$p$ = 1.00 | t(322) = -0.31<br>$p$ = 1.00 |
| *Useful Position > Upper Useful Position* | t(322) = 0.42<br>$p$ = 1.00 | t(322) = 0.89<br>$p$ = 1.00 | t(322) = 1.42<br>$p$ = 0.98 |
| *Useful Position > Lower Useful Position* | t(322) = 1.68<br>$p$ = 0.94 | t(322) = 0.28<br>$p$ = 1.00 | t(322) = -0.21<br>$p$ = 1.00 |
| *Useful Position > Saliency* | t(322) = 4.32<br>$p$ = 0.0019 | t(322) = 2.60<br>$p$ = 0.37 | t(322) = 3.01<br>$p$ = 0.15 |
| *Upper Useful Position > Lower Useful Position* | t(322) = 1.25<br>$p$ = 1.00 | t(322) = -0.61<br>$p$ = 1.00 | t(322) = -1.64<br>$p$ = 0.95 |
| *Upper Useful Position > Saliency* | t(322) = 3.90<br>$p$ = 0.0099 | t(322) = 1.71<br>$p$ = 0.93 | t(322) = 1.59<br>$p$ = 0.96 |
| *Lower Useful Position > Saliency* | t(322) = 2.64<br>$p$ = 0.35 | t(322) = 2.32<br>$p$ = 0.58 | t(322) = 3.23<br>$p$ = 0.087 |

**Table S1.** Results from post-hoc tests of the linear mixed models looking at theoretical RSMs in young adults. Tukey post-hoc tests from the linear mixed model investigating the effects of theoretical model, ROI and their interaction on the correlation coefficients between theoretical RSMs and neural RSMs in young adults only. We adjusted the *p*-value for a family of 15 estimates.

| | OPA | PPA | MPA |
|---|---|---|---|
| *Absolute Position > Useful Position* | $t(266) = -2.23$<br>$p = 0.62$ | $t(266) = -0.44$<br>$p = 1.00$ | $t(266) = -4.02$<br>$p = 0.0066$ |
| *Absolute Position > Upper Useful Position* | $t(266) = -1.15$<br>$p = 1.00$ | $t(266) = 0.086$<br>$p = 1.00$ | $t(266) = -2.90$<br>$p = 0.20$ |
| *Absolute Position > Lower Useful Position* | $t(266) = -1.38$<br>$p = 0.99$ | $t(266) = 0.098$<br>$p = 1.00$ | $t(266) = -2.84$<br>$p = 0.23$ |
| *Absolute Position > Saliency* | $t(266) = 0.93$<br>$p = 1.00$ | $t(266) = 0.92$<br>$p = 1.00$ | $t(266) = 0.46$<br>$p = 1.00$ |
| *Useful Position > Upper Useful Position* | $t(266) = 1.11$<br>$p = 1.00$ | $t(266) = 0.53$<br>$p = 1.00$ | $t(266) = 1.12$<br>$p = 1.00$ |
| *Useful Position > Lower Useful Position* | $t(266) = 0.88$<br>$p = 1.00$ | $t(266) = 0.54$<br>$p = 1.00$ | $t(266) = 1.17$<br>$p = 1.00$ |
| *Useful Position > Saliency* | $t(266) = 3.19$<br>$p = 0.098$ | $t(266) = 1.36$<br>$p = 0.99$ | $t(266) = 4.48$<br>$p = 0.0011$ |
| *Upper Useful Position > Lower Useful Position* | $t(266) = -0.23$<br>$p = 1.00$ | $t(266) = 0.012$<br>$p = 1.00$ | $t(266) = 0.053$<br>$p = 1.00$ |
| *Upper Useful Position > Saliency* | $t(266) = 2.08$<br>$p = 0.75$ | $t(266) = 0.83$<br>$p = 1.00$ | $t(266) = 3.36$<br>$p = 0.061$ |
| *Lower Useful Position > Saliency* | $t(266) = 2.31$<br>$p = 0.58$ | $t(266) = 0.82$<br>$p = 1.00$ | $t(266) = 3.31$<br>$p = 0.071$ |

**Table S2.** Results from post-hoc tests of the linear mixed models looking at theoretical RSMs in older adults. Tukey post-hoc tests from the linear mixed model investigating the effects of theoretical model, ROI and their interaction on the correlation coefficients between theoretical RSMs and neural RSMs in older adults only. We adjusted the *p*-value for a family of 15 estimates.

| | F-test | *p*-value | ES [95%-CI] |
|---|---|---|---|
| *Half* - DOWN x *Full* - DOWN **vs.** *Half* - UP x *Half* - DOWN | F(1, 778) = 12.26 | $p < 0.001$ | 0.016 [0.0030, 0.037] |
| *Half* - DOWN x *Full* - DOWN **vs.** *Half* - DOWN x *Full* - UP | F(1, 778) = 35.16 | $p < 0.001$ | 0.043 [0.020, 0.074] |
| *Half* - DOWN x *Full* - DOWN **vs.** *Half* - UP x *Full* - DOWN | F(1, 778) = 29.60 | $p < 0.001$ | 0.037 [0.015, 0.066] |
| *Half* - DOWN x *Full* - DOWN **vs.** *Full* - UP x *Full* - DOWN | F(1, 778) = 21.44 | $p < 0.001$ | 0.027 [0.0090, 0.053] |
| *Half* - DOWN x *Full* - DOWN **vs.** *Half* - UP x *Full* - UP | F(1, 778) = 0.16 | $p = 0.69$ | 0.00020 [0.00, 0.0067] |
| *Half* - UP x *Full* - UP **vs.** *Half* - UP x *Half* - DOWN | F(1, 778) = 9.66 | $p = 0.0020$ | 0.012 [0.0017, 0.032] |
| *Half* - UP x *Full* - UP **vs.** *Full* - UP x *Half* - DOWN | F(1, 778) = 30.64 | $p < 0.001$ | 0.038 [0.016, 0.068] |
| *Half* - UP x *Full* - UP **vs.** *Half* - UP x *Full* - DOWN | F(1, 778) = 25.47 | $p < 0.001$ | 0.032 [0.012, 0.060] |
| *Half* - UP x *Full* - UP **vs.** *Full* - UP x *Full* - DOWN | F(1, 778) = 17.94 | $p < 0.001$ | 0.023 [0.0065, 0.047] |
| *Full* - DOWN x *Full* - UP **vs.** *Half* - DOWN x *Half* - UP | F(1, 778) = 1.26 | $p = 0.26$ | 0.0016 [0.00, 0.012] |

**Table S3.** Results from post-hoc tests of the overall linear mixed model. The post-hoc F-tests for pairwise comparison from the overall linear mixed model investigating the effects of pairwise comparison, ROI and age group on the neural RSMs and neural RSMs. The effect size corresponds to partial eta squared $\eta_p^2$.
